# Robust summarization and inference in proteome-wide label-free quantification

**DOI:** 10.1101/668863

**Authors:** Adriaan Sticker, Ludger Goeminne, Lennart Martens, Lieven Clement

## Abstract

Label-Free Quantitative mass spectrometry based workflows for differential expression (DE) analysis of proteins impose important challenges on the data analysis due to peptide-specific effects and context dependent missingness of peptide intensities. Peptide-based workflows, like MSqRob, test for DE directly from peptide intensities and outperform summarization methods which first aggregate MS1 peptide intensities to protein intensities before DE analysis. However, these methods are computationally expensive, often hard to understand for the non-specialised end-user, and do not provide protein summaries, which are important for visualisation or downstream processing. In this work, we therefore evaluate state-of-the-art summarization strategies using a benchmark spike-in dataset and discuss why and when these fail compared to the state-of-the-art peptide based model, MSqRob. Based on this evaluation, we propose a novel summarization strategy, MSqRobSum, which estimates MSqRob’s model parameters in a two-stage procedure circumventing the drawbacks of peptide-based workflows. MSqRobSum maintains MSqRob’s superior performance, while providing useful protein expression summaries for plotting and downstream analysis. Summarising peptide to protein intensities considerably reduces the computational complexity, the memory footprint and the model complexity, and makes it easier to disseminate DE inferred on protein summaries. Moreover, MSqRobSum provides a highly modular analysis framework, which provides researchers with full flexibility to develop data analysis workflows tailored towards their specific applications.

## 1 Introduction

Label-Free quantitative (LFQ) mass spectrometry (MS) based workflows have become standard practice in quantitative proteomics (e.g. Goeminne et al. [2018], Tebbe et al. [2015]). This technology typically starts with protein extraction followed by an enzyme digestion step to produce shorter peptides. The thus obtained peptide mixture is then analyzed in a mass spectrometer where intact peptide masses and their intensities are measured, resulting in a so-called MS1 spectrum. In typical LFQ, the intensities of the thus recorded peaks are taken as proxies for peptide abundance. In order to identify the peaks observed in the MS1 spectrum, these peaks are first isolated in the instrument, and then subjected to fragmentation. Each of the resulting fragmentation spectra (so-called MS2 spectra) is then used for peptide identification. In LFQ, each sample is separately analyzed on the mass spectrometer, and differential expression is obtained by comparing relative intensities between runs for the same identified peptide. Goeminne et al. [2018]

However, this workflow also induces challenging data analysis problems. First, different peptides from the same protein often have very distinct physio-chemical properties, leading to large differences in their MS1 intensities even though these peptides are all equally abundant (Supplementary Figure 1 panel A1). Second, due to technological constraints not all peptides can be subjected to fragmetation. Indeed, only those peptides with the highest MS1 intensities within a certain retention window are typically selected for fragmentation. Tu et al. [2014] As a result, the identification in any given run depends not only on the abundance of that peptide, but also on the abundances of any co-eluting peptides. There can thus be context-depending missingness in a given run. Moreover, there are many other potential sources of (random or non-random) missingness, including peptide misidentification, ambiguous matching of MS1 peaks, and poor quality MS2 spectra. Lazar et al. [2016] Hence, there is considerable variation in terms of the peptides that are identified in each of the different MS runs in an experiment. Taken together, the identification issue and the peptide specific effects on quantification have a severe impact on the downstream summarization of peptide intensities towards protein abundances. Goeminne et al. [2015]

Indeed, because of these issues, simple summarization methods such as the mean or median peptide intensity are known to give unreliable protein abundance estimates Goeminne et al. [2015] and more advanced summarization strategies have therefore been proposed for LFQ data in the literature Silva et al. [2005], Cox et al. [2014], Zhang et al. [2018], Choi et al. [2014]. In Figure 1 panel A we show the performance of these different data analysis strategies on a benchmark dataset. Notably, we observe huge differences in performance between the different summarization strategies, which are driven by the absolute abundance, and any differences in this abundance, of a protein between conditions. Moreover, none of the summarization strategies outperforms the others across all conditions.

**Figure 1:**
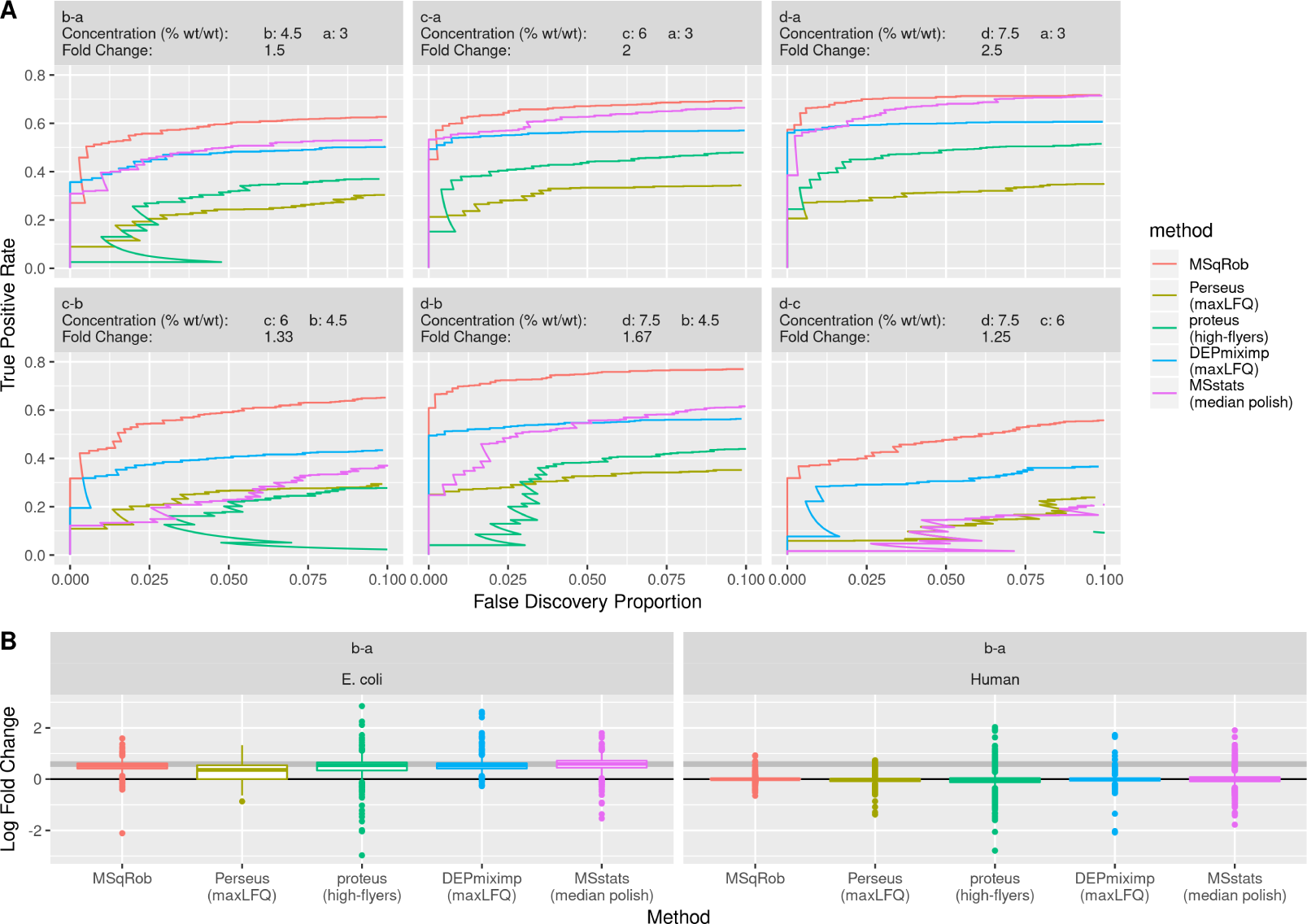
Comparison of current state-of-the-art tools for DE analysis of proteins. We compare one peptide based tool, MSqRob (red) and four summarization based tools: Perseus (green) and Differential Enrichment analysis of Proteomics data (DEP) with mixed imputation (blue) are both based on maxLFQ protein intensities. MSstats uses median polish summarised protein intensities (purple), while Proteus uses high-flyers summarization (green). The data consists of *E. Coli* proteins spiked at different concentrations (a, b, c and d) in a human proteome. The plot in Panel A shows the performance of each method for all pairwise comparisons. MSstats outperforms Proteus, DEP and Perseus at higher fold changes but drops in performance to Perseus levels at the lowest fold change. Proteus outper-forms Perseus at higher fold changes, but is less performant at the lowest fold change. MSqRob always outperforms the other methods. The box-plots in panel B show estimated log fold changes of differentially (*E. Coli*) and non-differentially (human) expressed proteins in the a vs b comparison. Perseus has biased fold changes for the *E. Coli* proteins, but has more precise fold changes for human proteins than DEP and MSstats. MSqRob has more precise and more acurrate fold changes than any other method.

In order to avoid these summarization issues, peptide-based models, such as MSqRob Goeminne et al. [2016], allow to test for differentially expression (DE) of proteins directly from the observed peptide intensities. The result is that these methods uniformly outperform summarization based methods (Figure 1 panel A). Indeed, by modeling peptide intensities directly, MSqRob naturally accounts for differences in peptide characteristics, and for differences in the number of identified peptides for a given protein in each sample, resulting in a bias reduction and a better uncertainty estimation on the fold change estimates. However, the MSqRob method also suffers from some drawbacks compared to summarization methods. MSqRob has to introduce random sample effects to account for correlation between the peptide intensities for a given protein in the same sample. This makes data analysis computationally more demanding, renders appropriate degrees of freedom of the test statistics unavailable, and even approximating these is impossible due to imbalances in the peptides across samples. The use of random effects also makes it difficult to disseminate the method towards non-specialised end-users as the interpretation of the result becomes correspondingly more complex. Moreover, MSqRob does not readily provide protein summaries for each sample, which are important for end-users to explore and visualise the data, and for further processing in downstream applications.

We therefore here introduce a novel estimation strategy for MSqRob using a two stage approach, which we call MSqRobSum. MSqRobSum provides robust protein level summaries that account for peptide specific effects, which are then further processed using robust ridge regression. Hence, MSqRobSum combines the advantage of MSqRob’s robust inference framework with the benefits of summarization, which allows fast and modular data analysis workflows. In addition, these workflows benefit from the straightforward visualization and interpretation of results at the protein level that is offered by MSqRobSum. We illustrate the high performance of MSqRobSum on a spike-in dataset and illuminate why it outcompetes existing state-of-the-art summarization-based tools for DE in LFQ MS-based quantitative proteomics.

## 2 Materials and methods

We performed a comparison of current state-of-the-art software tools for DE analysis of proteins on a benchmark spike-in dataset. We compared one peptide based tool, MSqRob and four summarization based tools: Proteus, Perseus, MSstats, and Differential Enrichment analysis of Proteomics data (DEP). For all tools we aimed to use the default workflow as suggested by the respective documentation. We also introduce our own novel summarization strategy for DE analysis, MSqRobSum, which aims to maintain MSqRob’s superior performance while also providing useful protein expression summaries.

### 2.1 Spike-in Dataset

The performance of MSqRobSum and other state-of-the-art software tools for differential expression analysis is benchmarked using a publicly available dataset (PRIDE identifier: PXD003881 Shen et al. [2018]). *E. Coli* lysates were spiked at five different concentrations (3%, 4.5%, 6%, 7.5% and 9% wt/wt) in a stable human background (four replicates per treatment). The twenty resulting samples were run on an Orbitrap Fusion mass spectrometer. Raw data files were processed with MaxQuant (version 1.6.1.0, Cox and Mann [2008]) using default search settings unless otherwise noted. Spectra were searched against the UniProtKB/SwissProt human and *E. Coli* reference proteome databases (07/06/2018), concatenated with the default Maxquant contaminant database. Carbamidomethylation of Cystein was set as a fixed modification, and oxidation of Methionine and acetylation of the protein amino-terminus were allowed as variable modifications. *In silico* cleavage was set to use trypsin/P, allowing two miscleavages. Match between runs was also enabled using default settings. The resulting peptide-to-spectrum matches (PSMs) were filtered by MaxQuant at 1% FDR. In all analyses, *E.coli* proteins are labeled as DE (true positives), and all human proteins as equally expressed (true negatives).

To benchmark performance and FDR control of these different quantification strategies, the False Discovery Proportion (FDP) and True Positive Rate (TPR) of a set of proteins returned by the method were calculated, with

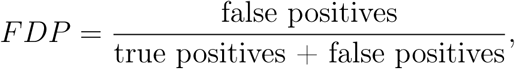

and,

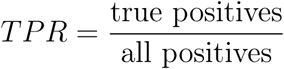

We define a set of significant DE proteins as the proteins with a p-value lower then a certain threshold. The FDP is then the fraction of human proteins in the set of human and *E. Coli* proteins recovered, while the TPR is the fraction of all *E. Coli* proteins recovered.

### 2.2 Proteus analysis

We performed the default workflow in the R package Proteus (0.2.9) starting from the PSM values as reported in the evidence.txt file in MaxQuant’s output.Gierlinski et al. [2018] Proteins that are only identified as contaminants or reversed sequences are removed from the data set. The intensities of PSMs in a given sample that can be assigned to the same peptide sequence are summed. Peptide intensities are summarised to protein intensities using the high-flyer method. Silva et al. [2005] Peptides were assigned to their leading razor protein. Protein intensities are normalized to the median, and median intensities in each sample are equal. Protein intensities are log_2_ transformed.

DE of proteins is analysed in Proteus with empirical Bayes moderated t-tests using the bioconductor limma package Ritchie et al. [2015]. Note, that we surpress an index for protein in all our model specifications for notational convenience.

In the limma analysis the following protein-wise linear models are considered:

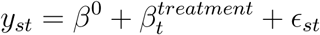

with *y*_*st*_ the normalised log_2_-transformed protein intensity in sample *s* of treatment *t, β*^0^ the intercept, 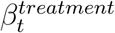 the effect of spike-in condition *t*, and, *ϵ*_*st*_ the protein-wise random error terms, which are assumed to be normally distributed with mean 0 and variance *σ*^2^. The variances *σ*^2^ are estimated with empirical Bayes, which stabilises the estimates by borrowing strength across proteins. Proteus corrects for multiple testing using the Benjamini-Hochberg FDR procedure.

### 2.3 Perseus analysis

We performed a standard Perseus workflow s tarting f rom t he MaxLFQ protein summaries calculated by MaxQuant. MaxLFQ protein summaries are normalized and summarised intensity values for each protein in a given sample. We can summarise the maxLFQ method as follows. The median ratio of the common peptides from a protein in all pairwise sample comparisons is calculated. Non-linear least-squares regression on these ratios is used to define a n o ptimal p rotein e xpression p rofile ac ross sa mples. Th is pr ofile is rescaled to match the total summed peptide intensities from this protein in all samples. Cox et al. [2014] MaxLFQ protein summaries, as reported in MaxQuant’s proteinGroups.txt file were f urther a ssessed i n Perseus version 1.6.0.7. Proteins that are only identified b y a m odification si te, contaminants, and reversed sequences are removed from the data set. Protein-wise two-sample t-tests on the log_2_ transformed maxLFQ values are performed for all pairwise treatment combinations. Perseus corrects for multiple testing using the Benjamini-Hochberg FDR procedure.

### 2.4 MSstats analysis

A standard MSstats (version 3.12 Choi et al. [2014]) workflow starts from the peptide intensities reported in MaxQuant’s evidence.txt file. Peptides with only one or two measurements across all samples, and peptides that occur in more then one protein are filtered out. When a peptide is measured multiple times in a sample, only the maximum intensity is kept. The log_2_ peptide intensities are median normalized and missing values are imputed using an Accelerated Failure Model (AFM). Peptide intensities are summarised to protein intensities using Tuckey’s median polish algorithm. Holder et al. [1979] MSstats build protein-wise linear models on these protein summaries.

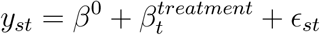

with *y*_*st*_ the normalised log_2_-transformed protein intensity in sample *s* of treatment *t, β*^0^ the intercept, 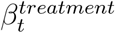 the effect of spike-in condition *t*, and, *ϵ*_*st*_ the protein-wise random error terms, which are assumed to be normally distributed with mean 0 and variance *σ*^2^. The multiple testing problem is corrected using the Benjamini-Hochberg FDR procedure.

### 2.5 Differential Enrichment analysis of Proteomics data (DEP)

MaxLFQ values are analysed with the standard workflow in the Bioconductor software package DEP version 1.2.0. Zhang et al. [2018] Proteins that are contaminants or that originate from reversed sequences are removed from the data set. Only proteins with no missing values in at least one treatment group are kept. The data are normalised using Variance Stabilizing Normalisation (VSN). von Heydebreck et al. [2002]

Missing values are imputed differently for proteins that are missing completely at random (MCAR), and proteins that are missing not at random (MNAR). Lazar et al. [2016] MCAR proteins are defined as proteins observed in at least one replicate for every condition, and these are imputed with k-nearest neighbors averaging. MNAR proteins are assumed to be missing under low abundance and are thus considered left-censored data. Proteins are labeled MNAR when completely missing in at least one condition and are imputed with a stochastic minimal value approach. In short, a value is drawn from a normal distribution centered around the first percentile of all observed protein expressions in the sample, and with a standard deviation estimated as the median protein-wise standard deviation.

DE of proteins is analysed in DEP with empirical Bayes moderated t-tests using the bioconductor limma package Ritchie et al. [2015], similar to the Proteus workflow. Multiple testing is corrected using an empirical FDR estimation approach as implemented in the R package fdrtool.

### 2.6 MSqRob analysis

The data is preprocessed using the MSnBase R/Bioconductor package version 2.6.2. Gatto and Lilley [2011] The analysis is done using the summarised peptide intensities as reported in the peptides.txt file in MaxQuant’s output. The data is normalised using Variance Stabilizing Normalisation (VSN). von Heydebreck et al. [2002] Proteins that are only identified by a modification site, contaminants, and reversed sequences are removed from the data set. To avoid ambiguity, peptide sequences attributed to both *E. coli* and human proteins are removed. Peptides that are only observed once across all samples are also removed. Finally, treatments in which a protein is only observed in one replicate are still included in the DE analysis for this protein.

MSqRob is a linear regression peptide-based mixed model. We consider the protein-wise models:

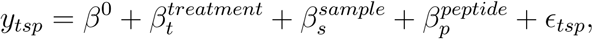

with *y*_*tsp*_ the normalised log_2_-transformed intensity of peptide *p* in sample *s* with treatment *t, β*^0^ the intercept, 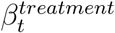 the effect of spike-in condition *t*, 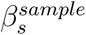a random effect that corrects for the correlation in measured expression levels between the peptides from the same protein in sample*s* (pseudo replication on the sample level), and, 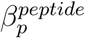 the effect of peptide *p*. Again the error term *ϵ*_*tsp*_ is assumed to be normally distributed with mean 0 and variance *σ*^2^.

When only one peptide is measured for a protein in all samples, the model reduces to:

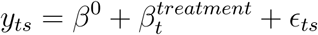

The parameters for *treatment* and *peptide* are tuned using penalised estimation by exploiting the link between random effects and ridge regression. Variability in the parameter estimators is reduced by shrinkage towards zero when there are only few observations. This protects against overfitting and makes the estimators more stable and accurate. The influence of outliers is weighed down by M-estimation using Huber weights. The variance of the protein-wise random error terms *ϵ*_*tsp*_ are again estimated with limma’s empirical Bayes variance estimator.

Multiple testing is corrected using the Benjamini-Hochberg FDR procedure.

### 2.7 MSqRobSum analysis

MSqRob’s mixed model can also be estimated through a two-stage regression analysis. Molenberghs and Verbeke [2000] Here we first summarise peptide intensities to the protein level and subsequently test for DE on these protein summaries.

The same preprocessing is used as for the MSqRob analysis described in section 2.6. In the first stage we aggregate all normalised peptide intensities of a protein using robust regression with M-estimation using Huber weights. We consider the protein-wise linear model:

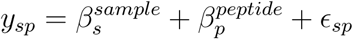

With *y*_*sp*_ the normalised log_2_-transformed intensity of peptide *p* in sample *s* and 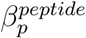 the effect of peptide *p*. By encoding the peptide effect as a sum contrast, 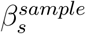can be interpreted as the mean intensity in sample *s* for this protein. The error term *ϵ*_*sp*_ is assumed to be normally distributed with mean 0 and variance 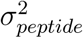.

In the second stage, we perform an MSqRob analysis on protein intensities with the reduced model:

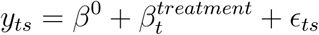

With *y*_*ts*_ the summarised log_2_-transformed protein intensity in sample *s* of treatment *t, β*^0^ the intercept, and, the effect of spike-in condition *t*. Again, the error term *ϵ*_*ts*_ is assumed to be normally distributed with mean 0 and variance *σ*^2^. We correct for multiple testing using the Benjamini-Hochberg FDR procedure.

We can expect a drop in performance in MSqRobSum compared to MSqrob because we lose information on the measured intensities, and introduce some random variation during summarization. We also don’t take into account that the covariance matrix of the estimated sample estimates is highly dependent on the number of measured peptide intensities in the sample Molenberghs and Verbeke [2000]. However, we expect that the resulting impact on performance is minimal in practice.

### 2.8 Software

Data preprocessing, statistical analysis and figures were done using the R programming language version 3.5.1. All R code is open sourced for reproducibility (https://github.com/statOmics/MSqRobSumPaper). The MSqRob algorithm has been implemented in R previously. Goeminne et al. [2016] However, we re-implemented MSqRob in R, and extended it to also allow for our proposed two-stage parameter estimation strategy, MSqRobSum. Because we fit a mixed model for each protein separately, we could easily parallelize the computations, which greatly speeds up the MSqRob and MSqRobSum analysis. In a full MSqRob peptide-level analysis, we typically allow for twenty iterations in the M-estimation using Huber weights for robust estimation of the model parameters. However, the MSqRob protein-level analysis in MSqRobSum only does one iteration by default. This sufficiently robustifies against outliers while maintaining proper FDR-control. The robust summarization, MSqrob and MSqrobSum algorithms are implemented as a open source R package *msqrobsum* (https://github.com/statOmics/MSqRobSum). The robust summarization algorithm is also ported to the *combineFeatures* function for summarization in the R bioconductor package *MSnbase*. Gatto and Lilley [2011]

## 3 Results

State-of-the-art methods and our novel MSqRob approach are all bench-marked using a dataset where an *E. Coli* proteome was spiked at five different concentrations in a human background. We first compare existing tools for DE analysis of LFQ based quantitative proteomics, and critically assess why the performance of summarization based approaches breaks down. Next, we show that our novel summarization based method, MSqRobSum, maintains the high performance of the peptide-level based approach MSqRob. We conclude this section by illustrating that MSqRobSum unlocks MSqRob towards modular data analysis workflows and we explain how and why MSqRobSum improves upon competitive summarization based approaches.

### 3.1 Comparison between methods

In this section we compare four summarization-based methods (Proteus, MaxQuant-Perseus, DEP, and MSstats), and one peptide-based model (MSqRob).

The Proteus workflow corrects for missingness of peptides under low abundance by summarizing using the high-flyer method, which provide protein-level intensities by taking the mean intensity of the three most intense peptides. Silva et al. [2005] Gierlinski et al. [2018]

However, this method does not correct for peptide specific effects and removes information by only using the top three peptide intensities. This introduces variability and bias in the estimated protein summaries (Supplementary Figure 1 panel B2). Indeed, the most abundant peptides typically differ between samples, leading to a low performance compared to all other methods (Figure 1 panel A).

The widely used MaxQuant-Perseus workflow is based on MaxQuant’s MaxLFQ summarization and subsequent statistical analysis with Perseus using t-tests (Cox et al. [2014]). MaxLFQ corrects for peptide specific effects by looking at pair-wise abundance ratios of shared peptides between samples. However, the heuristics in MaxLFQ often remove considerable information, which leads to increased missingness and imprecise summaries (Supplementary Figure 1 panel C3). In particular, comparisons that involve low spike-in concentrations often have too few shared peptides between samples, i.e. less than two, and these ratios are considered to be unreliable for summarization. Even though MaxLFQ corrects for peptide species by calculating ratios for shared peptides, it still appears to produces biased fold change estimates. The use of t-tests also results in suboptimal analysis as their variance estimator only includes the information of the data for the samples that are involved in the comparison. The summarization method combined with the less efficient downstream analysis often results in a low performance compared to the other methods (Figure 1).

The recent Differential Enrichment analysis of Proteomics data (DEP) software package greatly improved MaxLFQ based analysis by adopting a mixed imputation strategy for missing protein intensities that infers whether random missingness or missingness due to low abundance occurs Zhang et al. [2018]. It also provides a more robust downstream DE analysis using protein-wise linear models combined with empirical Bayes statistics (through the limma package Ritchie et al. [2015]). Hence, DEP produces both more accurate as well as more precise fold change estimates and vastly outperforms the Perseus analysis (figure 1).

MSstats Choi et al. [2014] is another popular software suite for proteomics data analysis. Initially, MSstats performed peptide-based modeling using linear mixed models. However, recent releases adopt summarization based workflows in which peptide intensities are first summarised to protein intensities, and linear modeling is then performed on the protein level. The default choice of summarization in MSstats is median polish, which corrects for peptide specific effects and is robust against outliers. Median polish is, however, unstable in the presence of too much missing data but this is alleviated in MSstats by imputing missing peptide intensities by default using an Accelerated Failure Model (AFM). However, unlike DEP’s mixed imputation strategy, AFM assumes that all intensities are missing because of low abundance, thus neglecting to take into account other sources of missingness.

The median polish summarization in MSstats produces more accurate fold change estimates compared to MaxLFQ (figure 1 panel B). However, while MSstats outperforms MaxLFQ based workflows at high fold changes (figure 1 panel A comparison b-a, c-a and d-a), its performance becomes increasingly worse in comparisons with low fold changes (figure 1 panel A c-b and d-c). This happens because the high fold change comparisons are achieved by a low concentration of spike-in proteins, with missingness predominantly caused by low abundance, while the low fold change comparisons contain a high concentration of spike-in proteins, with missingness often derived from other reasons. The missingness by low abundance assumption of MSstats is therefore much more likely to be violated for the low fold change, leading to a suboptimal ranking and a breakdown of MSstats for these comparisons. In contrast, DEP, which also accounts for random missingness, does not break-down for these comparisons.

It should be noted that DEP’s default preprocessing includes more stringent filtering for dubious proteins and thus returns less proteins overall than MSstats, which renders the better performance of MSstats in comparisons involving concentration a (b-a, c-a, d-a) superficial. Indeed, when only considering common proteins, DEP actually shows higher sensitivity then MSstats (Supplementary Figure 2 panel A).

MSqRob, finally, uses a peptide-based approach that provides robustness against outliers and overfitting by adopting M-estimation, ridge regression and a limma style empirical Bayes procedure for variance estimation Goeminne et al. [2016]. MSqRob thus derives unbiased fold change estimates with high precision and outperforms all summarization based models (figure 1 panel A). The increase in performance is even more apparent at low fold changes (figure 1 panel B comparison c-b and d-c).

### 3.2 MSqRobSum has similar overall performance to MSqRob

In this section, we show that we can fit the MSqRob model in a two-stage approach, with minimal impact on performance.

In the first stage of MSqRobSum we summarise peptide intensities in a sample to protein intensities using robust regression. This summarization is precise, and more robust than both high-flyer and maxLFQ summarization (Supplementary Figure 1). In the second stage, MSqRobSum provides precise and unbiased fold change estimates, comparable to MSqRob (Supplementary Figure 3).

MSqRobSum has a similar performance to MSqRob for medium to highly differentially expressed proteins (Figure 2 comparison b-a, c-a, d-a, and d-b). While the performance of MSqRobSum is lower than that of MSqRob for increasingly lower fold changes (Figure 2 comparison c-b and d-c), it should be noted that all summarization methods suffer from a drop in performance at lower fold changes (Figure 1).

**Figure 2:**
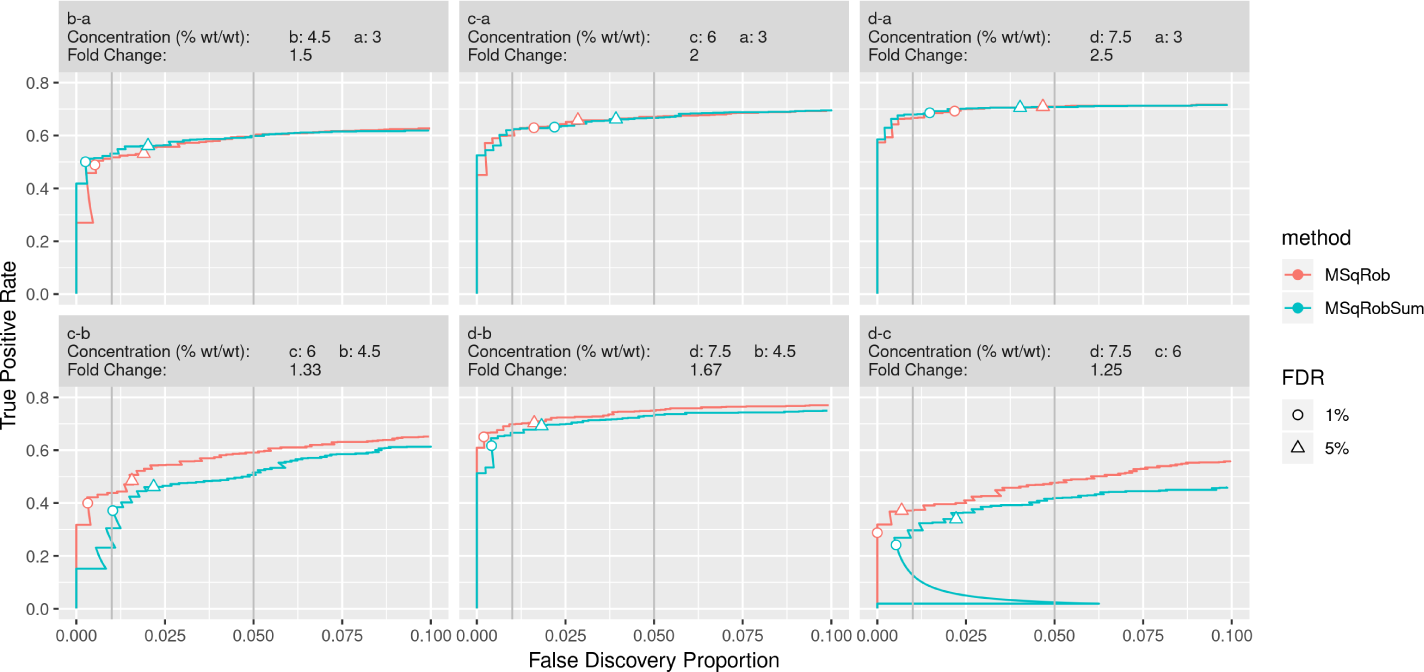
Comparison of performance in MSqRob and MSqRob-Sum. We compare the performance of MSqRob (red) and MSqRobSum (blue). The data consists of *E. Coli* proteins spiked at different concentrations (a, b, c and d) in a human proteome. The estimated 1% (circle) and 5% (triangle) FDR is controlled if it remains below 1% and 5% FDP, respectively (indicated by vertical grey lines). Performance of MSqRobSum is close to MSqRob in all comparisons, and MSqRobSum even outperforms MSqRob in the b-a comparison. The performance of MSqRobSum does decline a bit compared MSqRob at decreasing fold changes between treatments (eg. c-b and d-c), but the FDR is controlled in all comparisons except c-a and d-a, which both suffer from ion competition.

A major contributor to the performance drop of MSqRobSum is human protein Q9BZJ0, which has relatively low protein summaries for the samples in condition c due to outlying intensities of one peptide in all samples of condition c (Supplementary Figure 4). As a result, this protein receives a very low p-value from the MSqRobSum analysis for comparison d-c and is thus returned as a false positive at 1% FDR (Supplementary Figure 5). The MSqRob analysis, however, explicitely models the variance at the peptide level and the between sample variability and correctly rejects this protein.

The FDR is controlled at the 1% and 5% level for both MSqRob and MSqRobSum across almost the whole range of fold changes in differential expression (Figure 2), except in comparison c-a and d-a. The loss of FDR control in the latter comparison occurs because overspiking (high spike-in concentrations) causes increased ion competition between the peptides molecules in the sample Milac et al. [2012], Goeminne et al. [2015]. This in turn causes peptides with equal abundance in two samples to be less ionized in the sample with the higher total protein concentration, resulting in a lower measured intensity for those peptides in that sample.

This effect is clearly visible as the average estimated fold changes of the human proteins steadily decreases as the spiked-in *E. Coli* concentration increases (Supplementary Figure 6 A). At higher spiked-in *E. coli* concentrations, more human proteins thus appear to be differentially down regulated, and these additional false positives artificially inflate the estimated FDR (Supplementary Figure 6 B).

The rationale for switching to MSqRobSum instead of MSqRob is based on two issues with MSqRob. The first issue is that it is unclear which degrees of freedom should be used for the test with MSqRob. MSqrob uses the degrees of freedom of the variance at the peptide level (within sample variance), but these do not correspond to the degrees of freedom of the standard errors on the fold change estimates. Indeed, these standard errors include both the within sample variance and the between sample variance, and the correct degrees of freedom therefore vary between those of the within sample variance, and those that would be obtained for a tool that models the data at protein level. For unbalanced data, the correct degrees of freedom cannot be approximated and the results of MSqRob are thus bound to be too liberal.

The second issue is speed, as fitting the large mixed models in a peptide-level MSqRob workflow is computationally quite expensive. In contrast, the robust summarization in the first stage of MSqRobSum is a relatively cheap operation computationally. By switching to the two-stage approach in MSqRobSum, analysis time is reduced to less than a third of the MSqRob computation time. Parallelization of both methods maintains this speed difference, while decreasing processing time even further (Supplementary Figure 9).

### 3.3 MSqRobSum allows for a modular data analysis workflow

The MSqRobSum workflow consists of three steps: preprocessing, summarization with robust regression, and DE analysis with robust ridge regression. Because each step can have an important impact on the performance of the entire data analysis workflow, the decoupling of summarization and inference provides optimal flexibility to combine each of the MSqRobSum steps with other tools in modular workflows. To illustrate the usefulness of such modular workflows, we will start from the default Perseus workflow and we will show how each step in the MSqRobSum workflow ramps up the performance.

The default Perseus workflow consists of maxLFQ summarization combined with t-tests for statistical inference. Its performance is relatively low and the FDR is not controlled at either the 1% or the 5% level (Figure 3). However, exploratory data analysis revealed a strong batch effect across the samples which is undocumented in the experimental design. Batch effects should be corrected for during the statistical analysis but if undocumented, it is not always obvious if and how samples are organized in batches (Supplementary Figure 7). Often, normalization strategies already sufficiently correct for these sample effects. We can thus improve the performance and FDR control of the Perseus analysis by preprocessing the maxLFQ summarised intensities with VSN (Figure 3).

**Figure 3:**
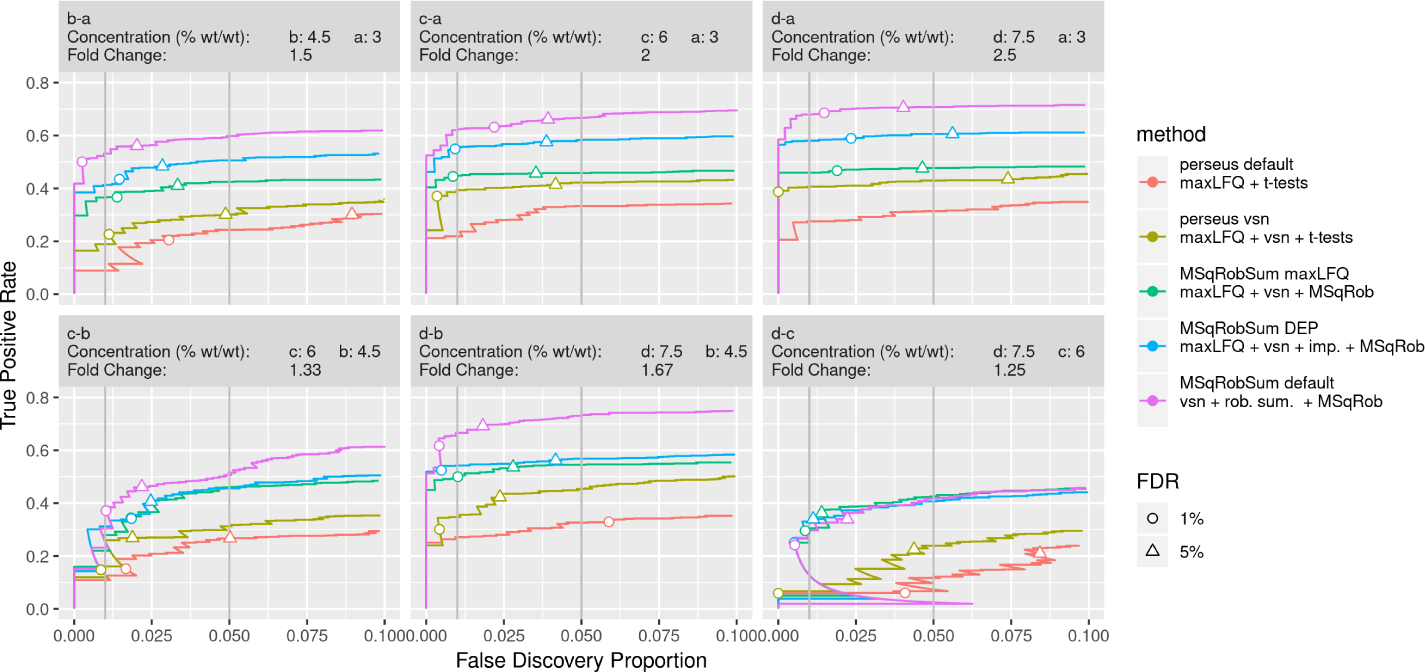
Improvements of DE analysis using a modular data analysis workflow. We show incremental improvements in DE analysis by incrementally changing components in the workflow. The data consists of *E. Coli* proteins spiked at different concentrations (a, b, c and d) in a human proteome. The circle and triangle are at 1% and 5% FDR, respectively, as estimated by the method. Perseus default performs t-tests on maxLFQ protein summaries for DE analysis. However the performance is low and FDR is not controlled. Adding VSN normalisation to the protein summaries boosts the performance of the DE analysis (perseus vsn). This workflow is further improved by swapping t-tests for MSqRob in the inference step (MSqRobSum maxLFQ). Adopting DEP’s mixed imputation scheme results in additional gain in perfomance (MSqRobSum DEP), and the best results are obtained by replacing maxLFQ and mixed imputation with our robust summarization (MSqRobSum default).

The statistical inference in Perseus is based on t-tests, which are underpowered when dealing with more than two conditions and other more complex study designs. We can therefore further improve performance by modeling intensities with MSqRob’s robust ridge regression approach, which allows for higher performance and good FDR control. Note, however, that FDR is not controlled in conditions c-a and d-a due to ion competition, as also highlighted above (Figure 3).

MaxLFQ’s summarization strategy, based on pairwise ratios between samples, is inefficient for samples with low concentrations, which leads to unstable summaries and/or missingness. DEP dealt with this through a context-dependent imputation strategy, which increases the power of the subsequent statistical inference (Figure 3). At high protein concentrations, there is low missingness and the effect of imputation will be small (Figure 3 comparison d-c).

With MSqRobSum, we correct for peptide-specific effects through a model-based robust summarization strategy which models the log-transformed peptide intensities directly through robust regression. This robust regression efficiently uses all available protein intensities and imputation such as used in MaxLFQ is therefore not required (Supplementary Figure 8). The full MSqRobSum workflow thus further boosts performance while maintaining good FDR control (Figure 3). Moreover, this MsqRobSum workflow uniformly outperforms all other modular approaches.

## 4 Discussion

In this work, we introduced MSqRobSum, a novel summarization-based method for LFQ which offers stable protein intensity estimation and highly performant protein DE analysis. We performed a benchmark study of different existing software implementations for summarization based LFQ methods and a state the state-of-the-art peptide based model, MSqRob. MSqRob uses the information on all peptides during statistical inference and outperforms all summarization based methods, which can only carry out inference on the protein summaries. However, MSqRob models are computationally quite expensive, can be hard to understand by experimentalists, include tests with unspecified degrees of freedom, and do not provide protein summaries for visualization and downstream processing. These MSqRob drawbacks are not present in summarization based methods. Indeed, summarization is usually a relatively cheap operation and reduces the number of data points, while the obtained protein summaries allow easy visual inspection of the data. The use of protein summaries also reduces model complexity and enables statistical inference with t-statistics that have well-defined degrees of freedom. However, many existing summarization-based methods suffer a considerable drop in performance compared to MSqRob (Figure 1). Our analysis shows that this drop in performance is dependent on issues with the summarization method used. Methods that do not take into account peptide specific effects, such as the high-flyer method in Proteus, show a clear drop in performance, while a method like MaxLFQ does consider peptide specific effects, but is based on heuristics and is not very data efficient. In MSqRobSum, we instead rely on robust regression for summarization, which allows to correct for peptide-specific effects, effectively exploits all data in its model based summarization, and is robust against outliers. Taken together, the result is a considerable boost in performance in the DE analysis when compared to MaxLFQ.

We also show that preprocessing is crucial for the performance of a DE workflow. The first type of such preprocessing is normalization, which can have a large impact on DE analysis (Figure 3). The second type of preprocessing is imputation of missing values, and this too can be beneficial (Figure 3). However, because several different imputation methods exist, and because each of these applies to different sources of missingness, best results are typically achieved when using a mixed imputation, where randomly missing values and values missing under low abundance are imputed differently Zhang et al. [2018]. It should be noted, however, that the robust modelling in MSqRobSum can safely omit imputation altogether (Supplementary Figure 8).

Another crucial component in LFQ is the statistical model for discovering DE proteins. Perseus utilises standard t-tests, but these are vastly underpowered compared to linear regression based models in more complex experimental designs. Moreover, MSqRob extends the linear model to robustify it against outliers and to improve uncertainty estimation Goeminne et al. [2015]. In the MSqRobSum workflow, we therefore use MSqRob’s robust linear model approach instead of t-tests on the protein summaries. This considerably improves performance of the DE analysis, reaching a level comparable to MSqRob for a wide range of DE proteins (Figure 3). And while MSqRob does show lower performance for increasingly lower fold changes in DE (Figure 2), all summarization methods suffer from a drop in performance in these cases, often more severe than that of MSqRobSum (Figure 3).

Moreover, the robust summarization approach has the merit that the entire analysis workflow has become modular: the provided robust protein abundance estimates can be used for visualisation and integration in other tools for DE, while MSqRob can now also start from protein summaries provide by other tools. This gives users considerable additional flexibility to develop modular workflows that are tailored towards their specific applications, and renders MSqRob future proof when novel and more performant summarization procedures become available.

## 5 Acknowledgments

We thank the students of the Statistical Genomics course, 2017-2018 Ghent University, who assisted us in assessing the initial implementation of MsqRob-Sum. L.M. is supported by European Union’s Horizon 2020 Programme under Grant Agreement 823839 [H2020-INFRAIA-2018-1] and Research Foundation Flanders (FWO) [grant number G042518N].

## 6 Supplementary material

**Figure 1:**
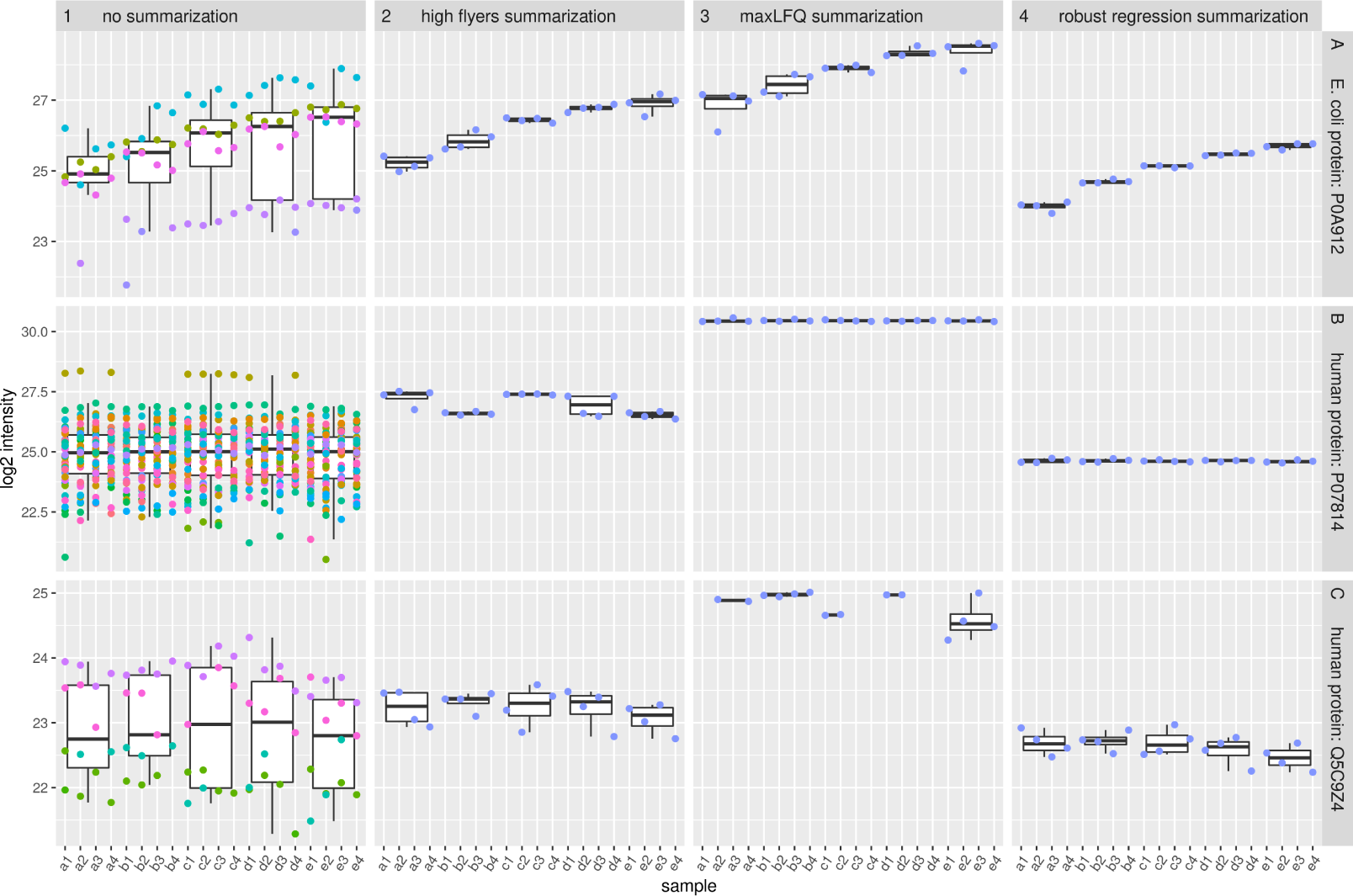
Comparison of different methods for estimating protein intensities. Rows A, B and C show the log_2_ intensities in each sample for one DE protein (POA912), and for two non-DE proteins (PO7814 and Q5C9Z4), respectively. Column 1 shows measured peptide intensities, with different peptides indicated by different colors. Columns 2, 3 and 4 show protein summaries estimated by high flyer, maxLFQ, and robust regression summarization, respectively. Plot A1 clearly illustrates peptide specific effects and context dependent missingness. Measured intensities from different peptides from the same protein show high variation and are prone to missingness for low abundant peptides (eg. purple peptide in samples a *versus* e). Robust regression provides stable protein intensity estimates that are more consistent for samples with the same concentration compared to high-flyer or maxLFQ summarization (eg. row A). high-flyer summaries are more unstable when high intensity peptides are missing in some samples (row B). MaxLFQ estimates are sometimes missing for proteins with few peptides (row C).

**Figure 2:**
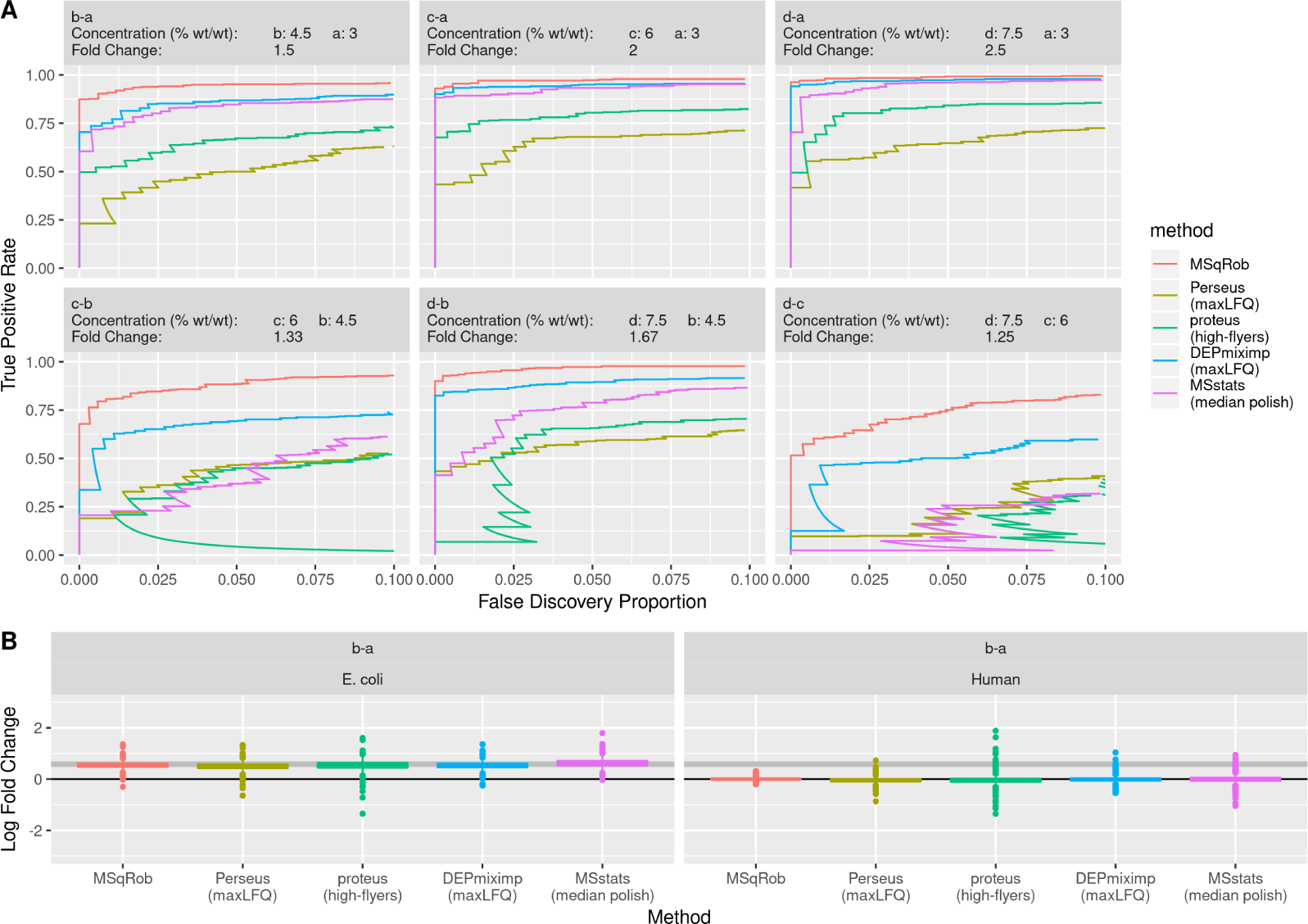
Comparison of current state-of-the-art tools for DE analysis of shared proteins. We compare one peptide based tool, MSqRob (red) and four summarization based tools: Perseus (green), and Differential Enrichment analysis of Proteomics data (DEP) with mixed imputation (blue), are both based on MaxLFQ protein intensities. MSstats uses median polish summarised protein intensities (purple). Proteus uses high-flyer summarization (green). The data consists of *E. Coli* proteins spiked at different concentrations (a, b, c and d) in a human proteome. the plot is based on proteins identified by all methods. The plots in Panel A show the performance of each method for all pairwise comparisons. MSstats outperforms Proteus, DEP and Perseus at higher fold changes, but its performance drops to Perseus levels at the lowest fold change. Proteus outperforms Perseus at higher fold changes, but is less performant at the lowest fold change. MSqRob always outperforms all other methods. The boxplots in panel B show estimated log fold changes of differentially (*E. Coli*) and non-differentially (human) expressed proteins in the a *versus* b comparison. Perseus has biased fold changes for the *E. Coli* proteins, but has more precise fold changes for human proteins than DEP and MSstats. MSqRob has more precise and more acurrate fold changes than any other method.

**Figure 3:**
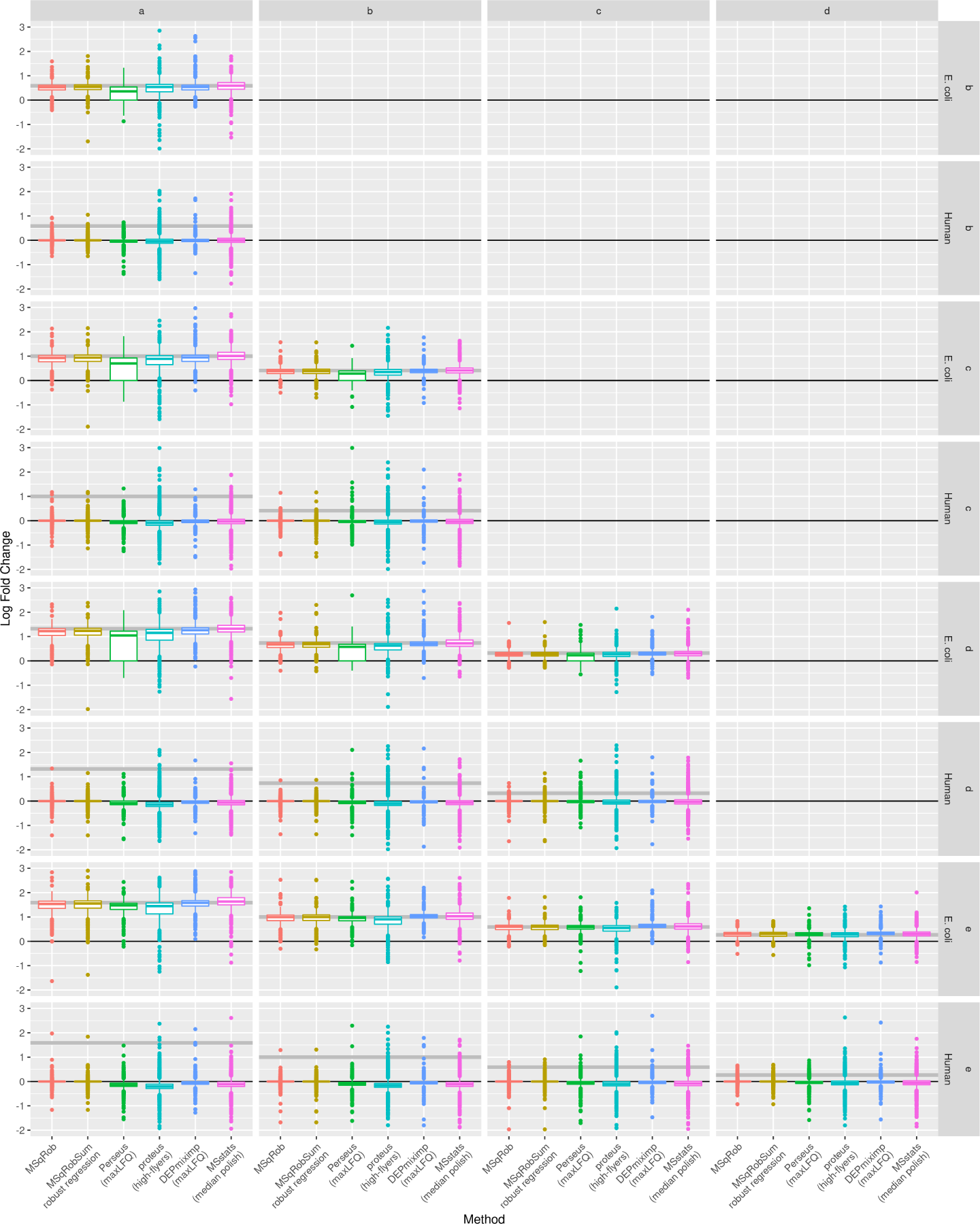
Comparison of current state-of-the-art tools for fold change estimation in proteins. We compare one peptide based tool, MSqRob (red) and four summarization based tools: Perseus (green), and Differential Enrichment analysis of Proteomics data (DEP) with mixed imputation (blue), are both based on MaxLFQ protein intenstities. MSstats uses median polish summarised protein intensities (purple). Proteus uses high-flyer summarization (green). The data consists of *E. Coli* proteins spiked at different concentrations (a, b, c, d and e) in a human proteome. Boxplots show estimated log fold changes of differentially (*E. Coli*) and non-differentially (human) expressed proteins. Perseus has biased fold changes for the *E. Coli* proteins, but has more precise fold changes for human proteins than DEP and MSstats. MSqRob and MSqRobSum both have more precise and more accurate fold changes than the other methods.

**Figure 4:**
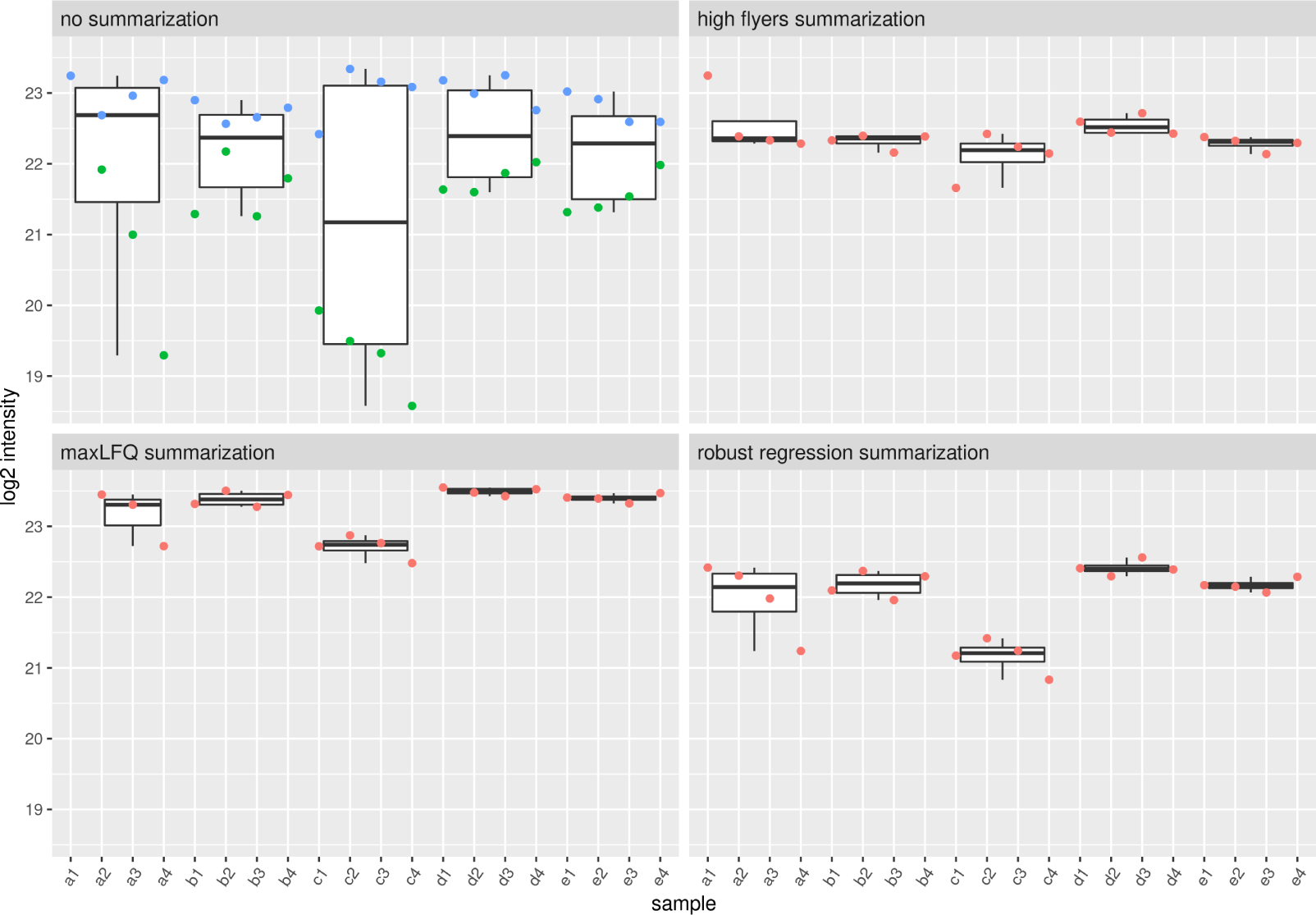
Peptide intensities and protein summaries for the human protein Q9BZJ0. The first panel shows the measured log2 intensities of the two peptides (green and blue) assigned to the human protein Q9BZJ0 for each sample. The other panel shows the protein summaries estimated by high-flyer, maxLFQ, and robust regression summarization, respectively. There is high variability on the log2 intensities of the green peptide between the samples and all four measurements in condition c are relatively low which results in a downward bias on the protein summaries. This lead to high uncertainty in the protein abundance estimates, especially for condition c. This uncertainty on the protein summaries, however, is not accounted for in summarization-based workflows. Peptide-based models like MSqRob explicitely model the variability at the peptide level as well as the between-sample variability and they are therefore less prone to flag this very specific edge case as a false positive.

**Figure 5:**
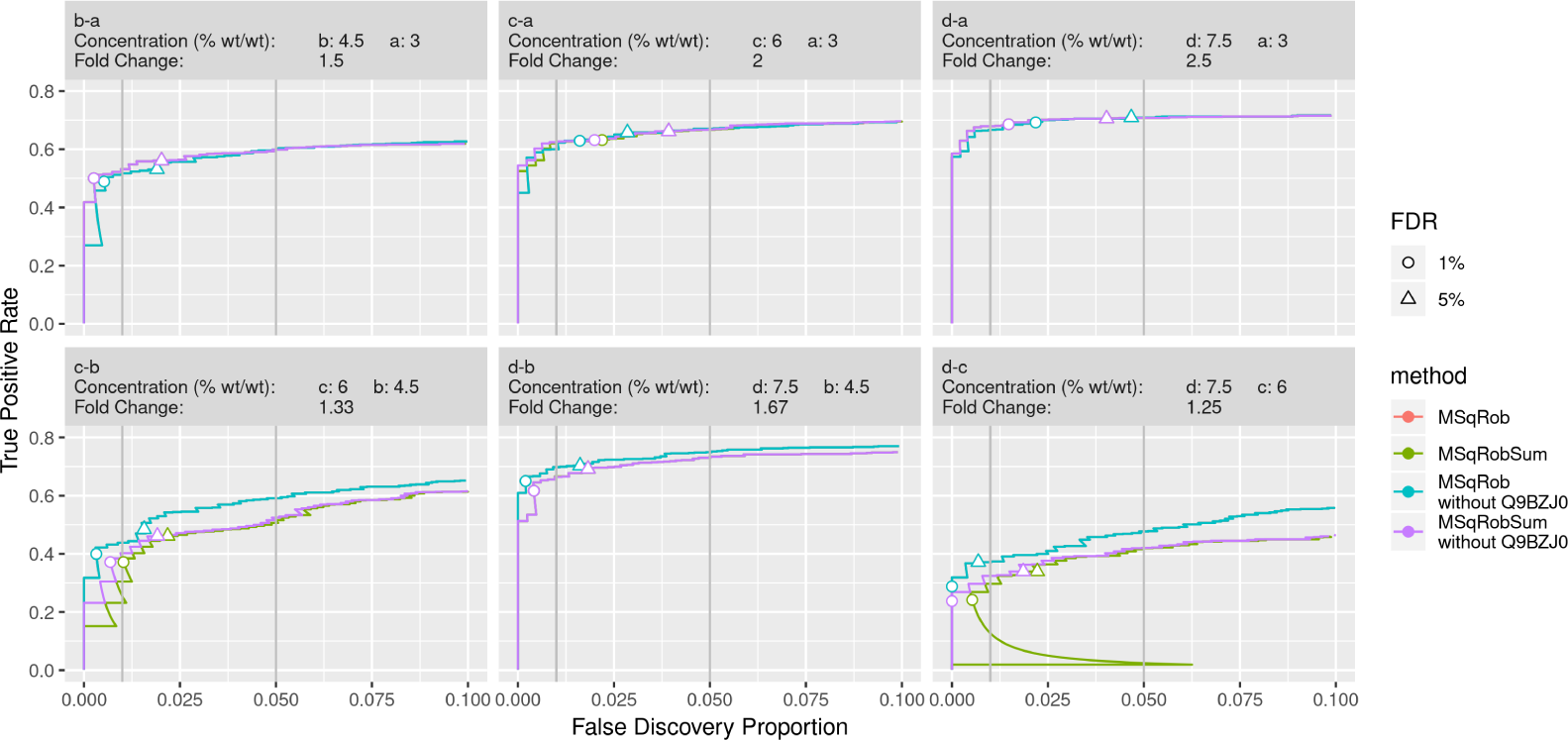
Performance comparison between MSqRob and MSqRobSum. We compare the performance of MSqRob and MSqRobSum on all proteins (respectively red and green) and all proteins except human protein Q5C9Z4 (respectively blue and purple). The data consists of *E. Coli* proteins spiked at different concentrations (a, b, c and d) in a human proteome. The estimated 1% (circle) and 5% (triangle) FDR is controlled if it remains below the 1% and the 5% FDP, respectively (vertical grey lines). Performance in MSqRobSum is close to MSqRob in all comparisons. MSqRobSum outperforms MSqRob in the b-a comparison. The performance of MSqRobSum is reduced compared to MSqRob at decreasing fold changes between treatments (eg. c-b and d-c). FDR is controlled in all comparisons except c-a and d-a due to ion competition. The sudden spike in the False discovery proportion in the d-c comparison can be explained by a low p-value for the human protein Q5C9Z4. Treatment c has relatively low protein summaries for Q5C9Z4 and removing this protein from the data results in an increased performance of MSqRobSum for all comparisons involving treatment c.

**Figure 6:**
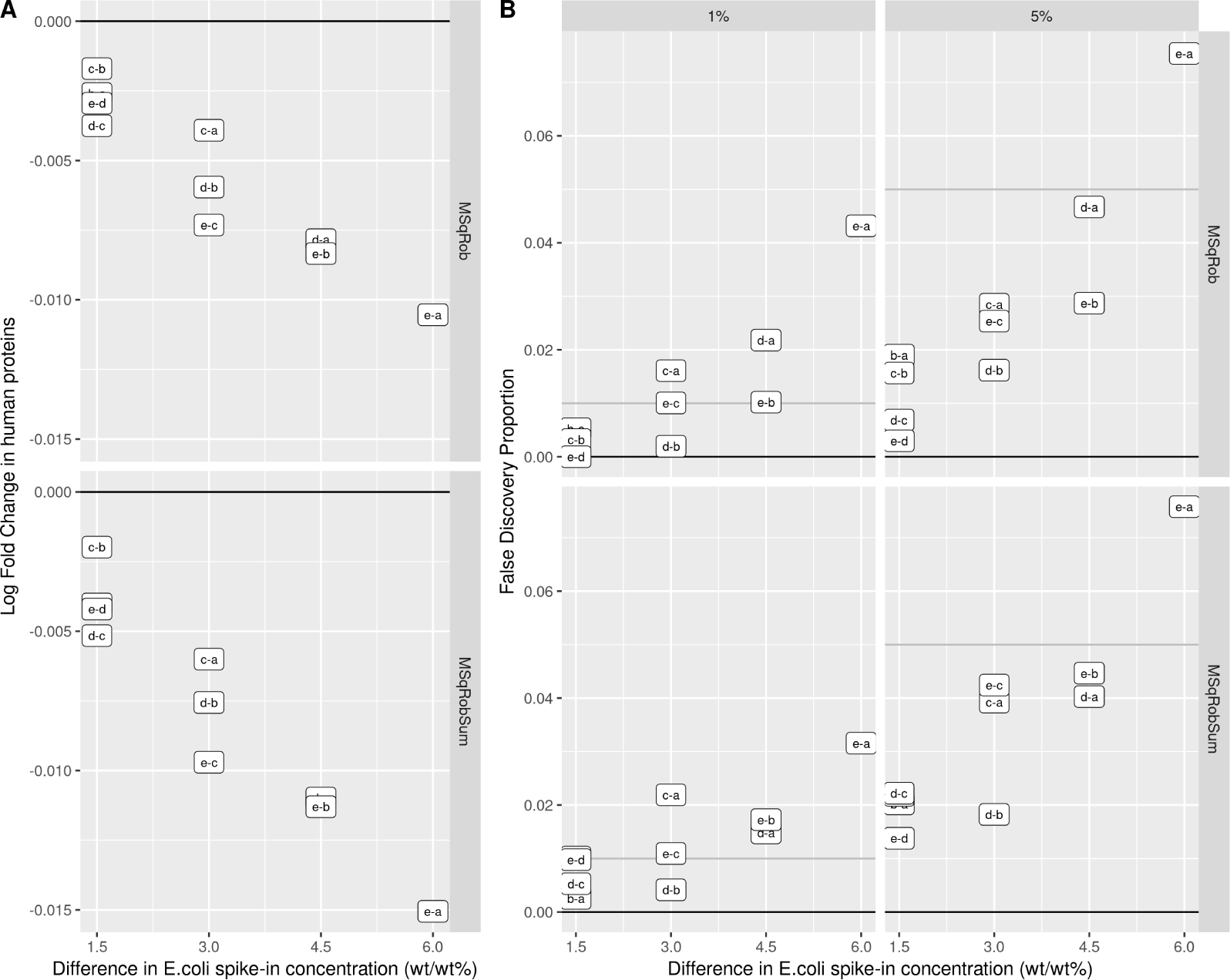
Overspiking can inflate the false discovery proportion. Panel A shows the false discovery proportion of a DE analysis controlled at 1% or 5% FDR for each comparison with different spike-in concentrations. At high *E. coli* spike-in concentration there is suppression of the MS1 intensities of the human proteins compared to low spike-in concentration due to ion competition. Panel A shows that the larger the difference in total protein concentration between samples, the larger the difference in estimated protein intensity of the non-DE human proteins, and the more likely these proteins appear down regulated. Panel B shows that the estimated FDR (grey line) is an underestimation of the true FDP (labelled boxes). Neither MSqRob nor MSqRobSum correctly controls the FDR for comparisons with high differences in spike-in concentration.

**Figure 7:**
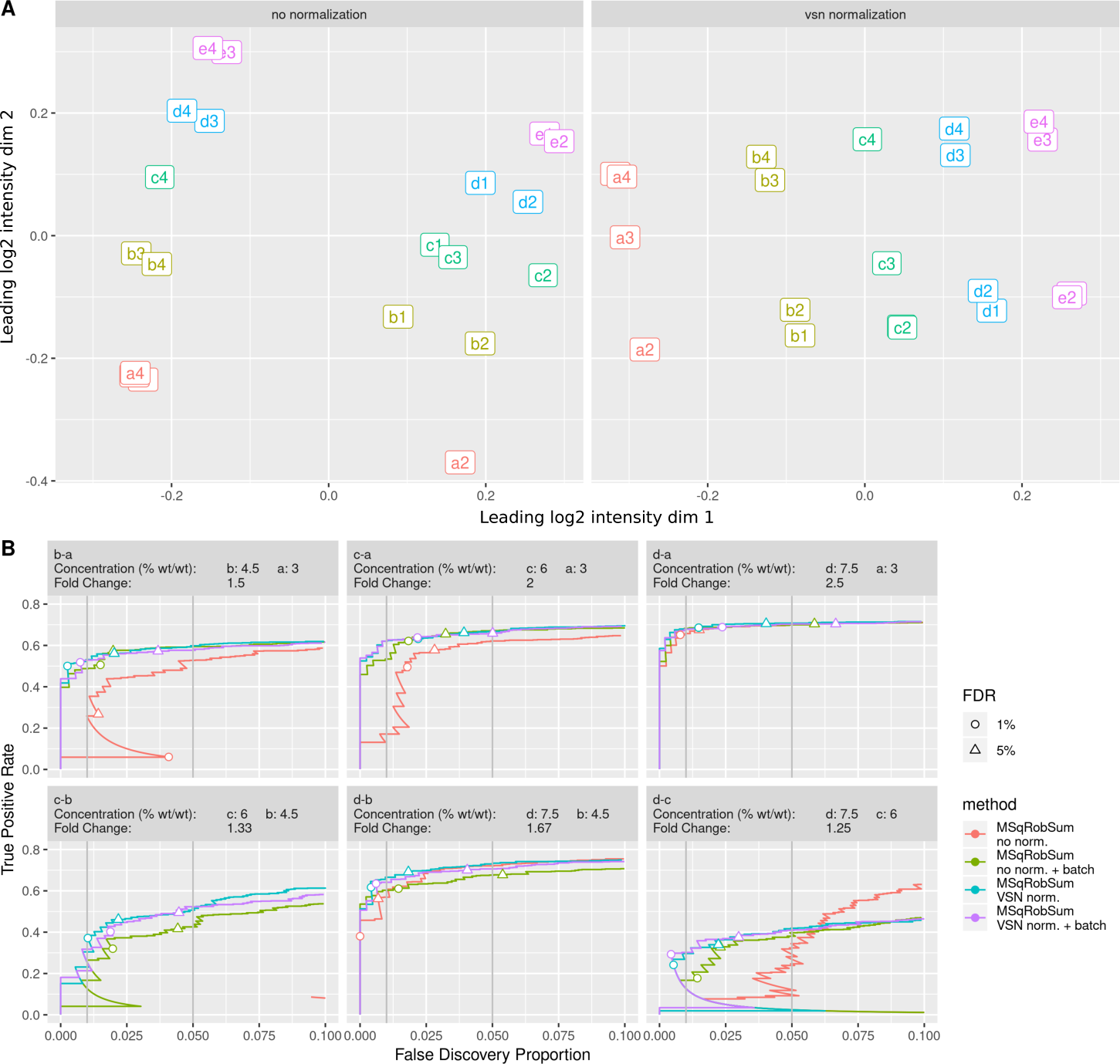
The effect of normalization on batch effects. Panel A shows MDS plots of the *log*_2_ sample intensities without normalization, and with VSN. Without normalization we see a clear separation of the samples in two groups along the first leading dimension. The spike-in condition effect (indicated by the 5 colours) only shows in the second dimension. After VSN, the unknown batch effect is still present but only in the second dimension; the condition effect now dominates the first dimension. Panel B shows the performance of MSqRobSum with and without normalization, and with and without inclusion of the batch effect into the model. Samples are assigned to two batches based on the MDS plot. Including the batch effect improves performance of the DE analysis considerably. The performance of the MSqRobSum analysis with VSN is similar to, or better than, the MSqRobSum model with the batch effect. Including a batch effect in the model after normalization does not improve performance further. The circle and triangle are at 1% and 5% FDR, respectively, as estimated by the method.

**Figure 8:**
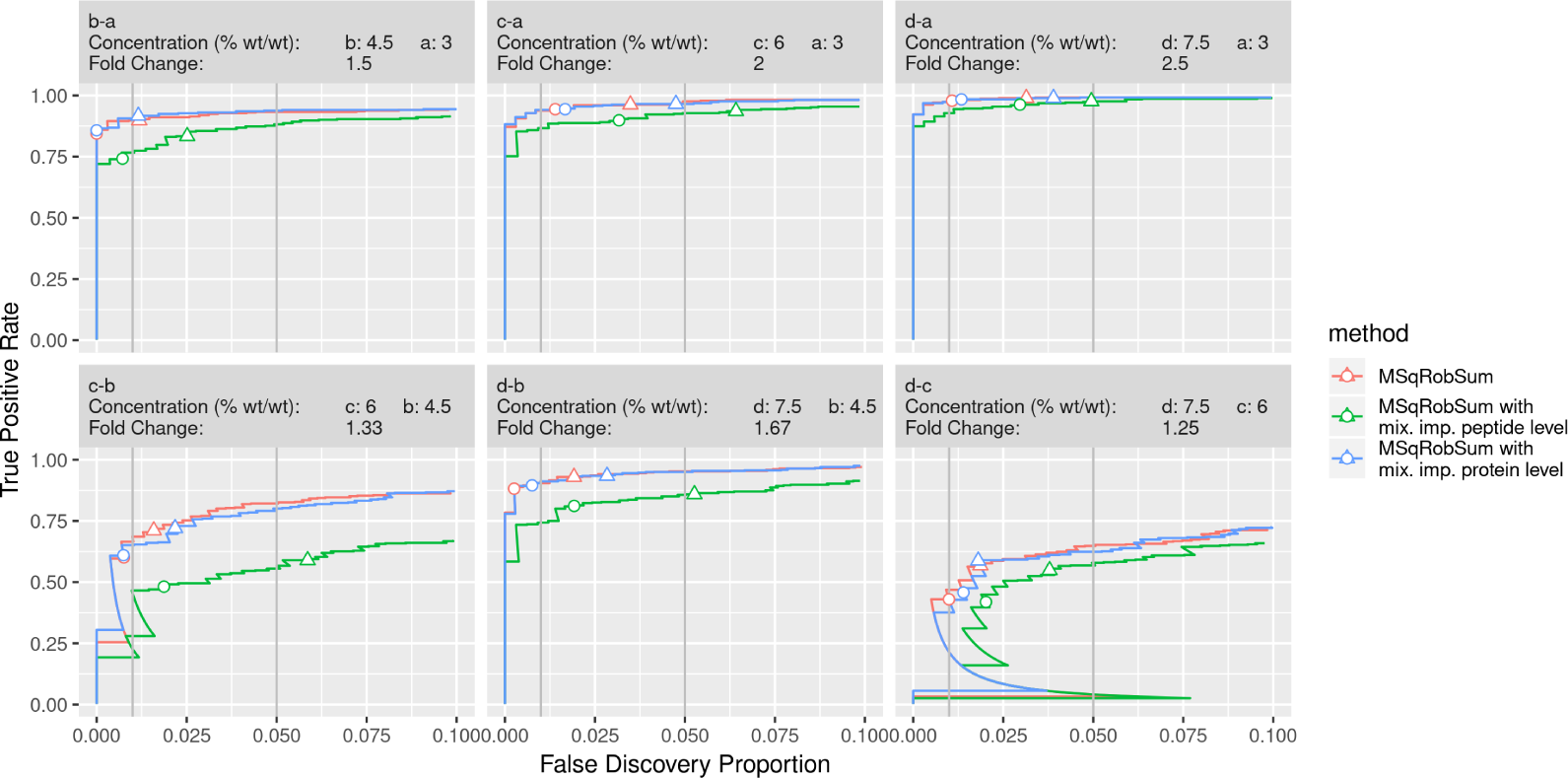
DEP’s mixed imputation does not improve MSqRobSum performance. The circle and triangle are at 1% and 5% FDR, respectively, as estimated by the method. Imputing the missing values in the protein summaries after robust summarization with DEP’s mixed imputation strategy does not improve the performance of MSqRobSum compared to the no imputation approach. Imputing missing values in the measured peptide intensities before robust regression actually degrades the performance of the DE analysis, and also often leads to loss of FDR control.

**Figure 9:**
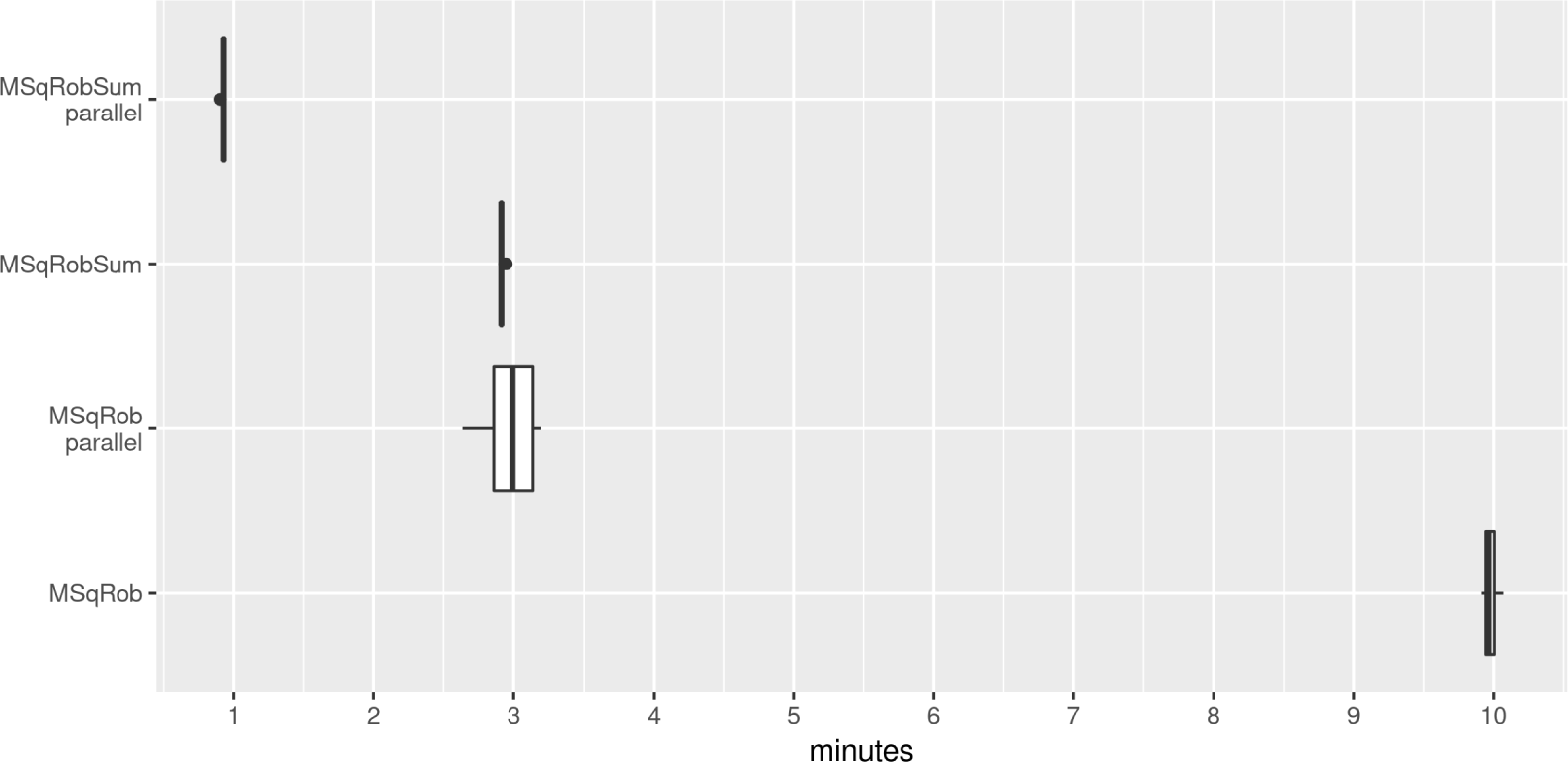
Speed comparison between MSqRob and MSqRobSum. All analyses were performed on the *E.Coli* spike-in benchmark data set. The same preprocessing is applied in all analyses and this preprocessing is excluded from the time measurements. Every analysis is run twenty times on a DELL Latitude laptop with eight i7-7820HQ CPU cores at 2.9 GHz and 31.3 GiB of RAM memory. The parallelized versions were allowed to use all eight available cores.

